# Sex differences in low-dose ethanol effects on motivated behavior and limbic corticostriatal activity

**DOI:** 10.1101/2025.10.07.680977

**Authors:** Christina M Curran-Alfaro, Kathleen G Bryant, Toni-Shae Ledgister, Sana Amin, Jacqueline M Barker

## Abstract

**Background:** Even at lower doses, ethanol exposure impacts both the brain and behavior. Emerging work has shown that chronic exposure to lower doses of ethanol may lead to inflexible behaviors and promote aberrant reward seeking. This study investigated the impact of chronic, low-dose ethanol exposure on neural substrates of reward and on motivated behavior.

**Methods:** Adult C57BL/6J mice were trained to self-administer sucrose. Throughout training, mice received an injection of low-dose ethanol (0.5g/kg) or saline, 1 hour after each session. Mice did not receive ethanol during testing. Mice were then tested in a PR task in which the reward magnitude of reinforcer was reduced (small or large) or increased (small or large). A subset of mice expressed a retrograde tracer in the nucleus accumbens (NAc), and cFos expression within NAc circuits was analyzed following a sucrose self-administration session.

**Results:** Chronic low-dose ethanol exposure altered behavioral responding in female mice following small changes in reward magnitude. Female mice showed divergent response patterns when there was a small reduction in reward magnitude, with greater proportions of ethanol-exposed female mice either increasing or decreasing responding versus controls. Following a small increase, low-dose ethanol female mice significantly increased responding versus controls. Female – but not male - mice exposed to chronic low-dose ethanol shifted behavioral strategy with a reduction in magazine ‘checking’ behavior. Low-dose ethanol exposure altered cFos expression within the prelimbic cortex and its projections to the NAc during reward seeking.

**Conclusions:** Chronic, low-dose ethanol altered behavioral responding and strategy in female mice in response to changes in reward value. Low-dose ethanol exposure impacted cFos induction in prelimbic cortex and its projections to NAc in both female and male mice. Future studies should investigate the consequences of chronic, low-dose ethanol on the brain and behavior to understand what underlying processes drive aberrant reward-seeking behaviors.

## Introduction

It is increasingly appreciated that even low doses of ethanol can impact health. This includes recognition from the World Health Organization that there is no safe amount of ethanol consumption as any level of ethanol consumption increases risk of negative health outcomes (Anderson et al., 2023). Despite this, consumption of ethanol is very common (WHO, 2024), necessitating a greater characterization of its impacts on brain and behavior. Low-dose ethanol exposure has distinct physiological effects from higher dose ethanol exposure. At lower doses, ethanol elicits a more stimulatory response, promoting disinhibition and relaxation, whereas at higher doses ethanol exposure induces sedative effects resulting in impaired motor function, coma and at severe levels, potentially death (Cui and Koob, 2017). Chronic exposure to lower doses of ethanol impacts both brain and behavior (Cui and Koob, 2017; Davidson et al., 1997; Daviet et al., 2022; Weafer and Fillmore, 2016). These neurobehavioral changes arising from chronic low-dose ethanol exposure contribute to increased risk for the development of alcohol use disorder (AUD) or other disorders associated with aberrant motivated behavior. Our previous work has shown that chronic, low-dose ethanol exposure enhanced motivation for a reward in a sex-specific manner (Bryant et al., 2022). However, it is not clear which aspects of motivated behavior and associated neurocircuitry are impacted by chronic low-dose ethanol exposure. Corticolimbic brain regions such as the prefrontal cortex (PFC) and basolateral amygdala (BLA) play a large role in cognitive processes necessary for flexible behavior and are known to be sensitive to higher doses of ethanol (Abernathy et al., 2010; Gass and Chandler, 2013; Klenowski et al., 2021; Koob and Volkow, 2016; Nippert et al., 2024; Pleil et al., 2015; Shields et al., 2022; Trantham-Davidson and Chandler, 2015). To expand on these findings, the impact of chronic low-dose ethanol exposure on behavioral sensitivity to changes in nondrug reward value was investigated using a progressive ratio task in which reward magnitude was either decreased or increased. The effect of chronic, low-dose ethanol on cFos expression within projections to the nucleus accumbens (NAc) was assessed following a sucrose self-administration session to determine the impact of low-dose ethanol on NAc circuit engagement. These findings give insight to chronic low-dose ethanol exposure effects on aberrant reward motivation and seeking.

## Materials and Methods

### Subjects

Adult male and female C57BL/6J mice from Jackson Laboratory (delivered at 9 weeks in age; *N* = 71 (35 females, 36 males; 1 female excluded due to failure to acquire the task) were used in the following studies. All studies were performed in accordance with approved protocols from the Drexel University Institutional Animal Care and Use Committee. All mice were housed in a vivarium with a standard 12h:12h light-dark cycle. Mice were acclimated to the facilities for at least one week before beginning experiments. Prior to the start of experiments, mice were food restricted to approximately 90% of their *ad libitum* weight, which lasted for the duration of the behavioral studies.

### Surgeries

In order to investigate the effects of low-dose ethanol on NAc circuitry, 24 adult C57BL/6J mice (*N* = 12 males and 12 females) underwent stereotaxic surgery for microinjections of a retrograde GFP tagged viral tracer (pAAV-hSyn-EGFP; Addgene, Watertown, MA; #50465) into the NAc. Surgeries were performed using isoflurane anesthesia. Virus was injected (0.2 uL infusion at 0.1 uL/minute) targeting 1.5 mm anterior/posterior (A/P), + 0.6 mm medial/lateral (M/L) and −4.7 mm dorsal/ventral (D/V) to bregma (0 mm). The syringe was left in place for 5 minutes for diffusion.

### Behavior training

All behavioral experiments were conducted using standard Med-Associates (St. Albans, VT) operant boxes. On one wall of each box there were two retractable levers on either side of a reward magazine containing a slot for pellet or liquid reinforcer delivery. Cue lights were directly above each lever but were not used in these experiments. On the opposite wall of each box, there were five nose pokes with lights which were not used for any of the experiments. The back wall, ceiling and door of the box were comprised of Plexiglas. All operant boxes were housed within sound attenuating chambers which had ventilation by fan and white noise.

Mice underwent operant training to self-administer 10% sucrose (20 uL per reinforcer delivery) on a fixed ratio 1 (FR1) schedule on two levers, where each lever press resulted in one reinforcer delivery. Levers were presented consecutively, and mice had access to each lever for 15 minutes, with the order of lever presentation counterbalanced across days for each animal. After acquisition of stable responding on the FR1 schedule (> 15 presses on each lever for 3 consecutive days), the reinforcement schedules were transitioned. One lever was assigned to be reinforced on a random interval (RI) schedule, which has been shown to promote automated behaviors, and the other on a variable ratio (VR) schedule, which promotes maintenance of goal-directed responding (Barker et al., 2018; Bryant et al., 2022; Dickinson et al., 2002; Gremel and Costa, 2013). The RI lever was reinforced at RI30 (the first lever press after, on average, 30 seconds is reinforced) for 3 days, followed by RI60 for 3 days. The VR lever was reinforced at VR5 (on average, the 5th lever press was reinforced) for 3 days followed by 3 days on a VR8 schedule. All progressive ratio testing was conducted on the lever trained using the RI schedule as behavioral impacts of training on this schedule on reward motivation have been observed following chronic, low-dose ethanol exposure (Bryant et al., 2022).

### Low-dose ethanol exposure

Mice were matched by sex to receive low-dose ethanol (0.5 g/kg, i.p.) or saline vehicle injections 1 hour after the initiation of each training session with a range of 13-39 days of training/exposure per animal. The 1-hour time point was chosen to match previous literature looking at the effects of low-dose ethanol exposure (Bryant et al., 2023, 2022). After training was completed, mice no longer received ethanol or saline injections and thus went into progressive ratio (PR) testing with a history of low-dose ethanol exposure.

### Progressive ratio testing

In order to assess the impact of chronic low-dose ethanol exposure on motivated behavior under changing reward conditions, 47 mice (24 males, 23 females) with a history of low-dose ethanol exposure (or vehicle controls) underwent PR4 testing. In these sessions, only the lever trained on the RI schedule was protracted. During the PR test, reinforcer delivery requirements progressively increased, with the number of lever presses required for each successive reward progressing in increments of 4 (1, 5, 9, 13, 17…). During training, reinforcer magnitude was 20 uL of 10% sucrose. To assess the effects of a history of low-dose ethanol exposure on sensitivity to changes in reward value - including increases vs. decreases in magnitude, and small vs. large changes - reinforcer magnitude was altered across PR testing sessions. PR testing was counterbalanced to assess small changes (from 20 uL to 15 or 25 uL) or large changes (from 20 uL to 5 or 30 uL) from the baseline reinforcer volume. The selected changes in reward volumes were consistent with reported detectable changes in reward volumes by mice (Nachev et. al, 2021). All testing was performed on the PR4 schedule and conducted across an 8-hour session until animals reached a “breakpoint” where they discontinued pressing (no lever presses for 5 min) (Bryant et al., 2022; Gourley et al., 2007).

### Immunofluorescence

To probe the effects of chronic low-dose ethanol exposure on NAc circuits involved with reward motivation and seeking, tissue generated from animals from a previously published behavioral data set (Bryant et al., 2022, 2024) were utilized. Following the completion of PR4 testing, animals with a retrograde tracer in the NAc underwent a 15-minute RI60 session (*N* = 12 males and 12 females). To detect cFos protein expression, mice underwent transcardial perfusion 90 minutes from the midpoint of the 15-minute session. Brains were postfixed in 4% paraformaldehyde solution then transferred to 30% sucrose for cryoprotecting. Brain tissue was then cut at 40 microns in preparation for immunofluorescence staining. Tissue was incubated in cFos rabbit (Abcam, Cambridge, UK; #ab222699, 1:500 dilution) and GFP goat (Abcam, Cambridge, UK; #ab5450, 1:1000 dilution) primaries in 4°C overnight. Tissue was then incubated in Alexa Fluor 594 anti-rabbit (Invitrogen, Carlsbad, CA; #A32754, 1:250 dilution) and Alexa Fluor 488 anti-goat (Jackson ImmunoResearch Labs, West Grove, PA; #705-545-147, 1:250 dilution) secondary antibodies for 2 hours at room temperature. Slices were then sorted and mounted onto slides for immunofluorescence microscopy imaging using a Nikon motorized microscope with 10x objective. Quantification of cFos, GFP and colocalization puncta analysis was conducted by a scientist blind to all groups/conditions using ImageJ and normalized by area of the region of interest. Labeled cells were quantified from slices at bregma −.95, −1.06, −1.23 mm for BLA, +1.97, + 1.69, +1.53 mm for PFC, and +1.7, +1.4, and +1.1 mm for NAc (9 slices/mouse; 3 slices/region) using the Franklin & Paxinos ‘The Mouse Brain Atlas’, 3^rd^ edition as reference.

### Statistical analysis

All statistical analyses were conducted using GraphPad Prism. Differences in group means were determined using unpaired t-tests or, for repeated behavioral testing, rmANOVA. To characterize population differences in behavior, responding during the PR4 test sessions was calculated as percent change. Using a quartile split, the mice that were in the highest 25% were identified as ‘increasers’, the bottom 25% as ‘decreasers’, while the middle 50% were considered not to change their behavior. A chi square was performed to analyze distribution of changes in breakpoint for each PR4 test (5uL, 15uL, 25uL and 35uL). ‘Checking’ behavior analysis was calculated for the complete PR session as well as for each block within the PR session as effort requirement for reinforcer delivery increased for each PR block using MATLAB (version R2022a; <u>Barker Lab Github</u>; https://github.com/bryantkg/Barker-lab-code).

## Results

### Low-dose ethanol does not impact sucrose self-administration acquisition

To determine if low-dose ethanol impacted sucrose self-administration, we compared lever responding in saline versus ethanol exposed mice. There was no effect of low-dose ethanol on lever responding during acquisition of sucrose self-administration [3way ANOVA: EtOH: F(1,43) = 0.6323, p = 0.4309]. There was a main effect of sex on lever responses during training [Sex: F (1, 43) = 5.846, p = 0.0199] as well as a main effect of training days [F (5.227, 224.7) = 12.07, p < 0.0001] with elevated responding compared to day 1 on days 4 - 9 [p’s < 0.05; Sidak’s] in both female and males. There were no interactions between training days, sex or EtOH during acquisition of sucrose self-administration [*Training days x EtOH*: F(5.227,224.7) = 0.7216, p = 0.6138; *Training days x Sex*: F(5.227,224.7) = 0.7509, p = 0.5918; *EtOH x Sex*: F(1, 43) = 2.100, p = 0.1545; *Training days x EtOH x Sex*: F(5.227,224.7) = 0.6860, p = 0.6409]. Due to this main effect of sex on responding during self-administration for basal magnitude of 20uL of 10% sucrose which may reflect differing baseline values of the reward and impact our interpretation regarding behavioral response to reward magnitude change or motivated behavior, we have segregated further behavior analysis within sex.

To assess the effects of chronic low-dose ethanol exposure on sucrose self-administration acquisition (FR1) [See behavioral timeline, **Fig 1A**], the number of days required to meet response criteria during training was compared in saline versus low-dose ethanol-exposed mice. Ethanol did not impact days required to reach acquisition criteria in female [unpaired t-test, t = 1.524, df = 21; p = 0.1424; **Fig 1B**] or male mice [unpaired t-test: t = 0.4683, df = 22, p = 0.6442; **Fig 1C**]. Two-way ANOVA indicated no effect of low-dose ethanol on lever responding across day in female [F(8,168) = 0.3668, p = 0.3933; **Fig 1D**] or male mice [F(8,176) = 1.030, p = 0.4154; **Fig 1E**]. A main effect of training days was observed in females [F(3.147, 66.08) = 5.189, p = 0.0024, Greenhouse-Geisser corrected], with elevated responding compared to day 1 on days 7, 8, and 9 [p’s < 0.05 Sidak’s]. A similar main effect of day was observed in males [F(5.645, 124.2) = 7.607, p < 0.0001, Greenhouse-Geisser corrected], with elevated responding vs day 1 on days 4, 6, 7, 8, and 9 [all p’s < 0.05, Sidak’s].

**Figure 1.**
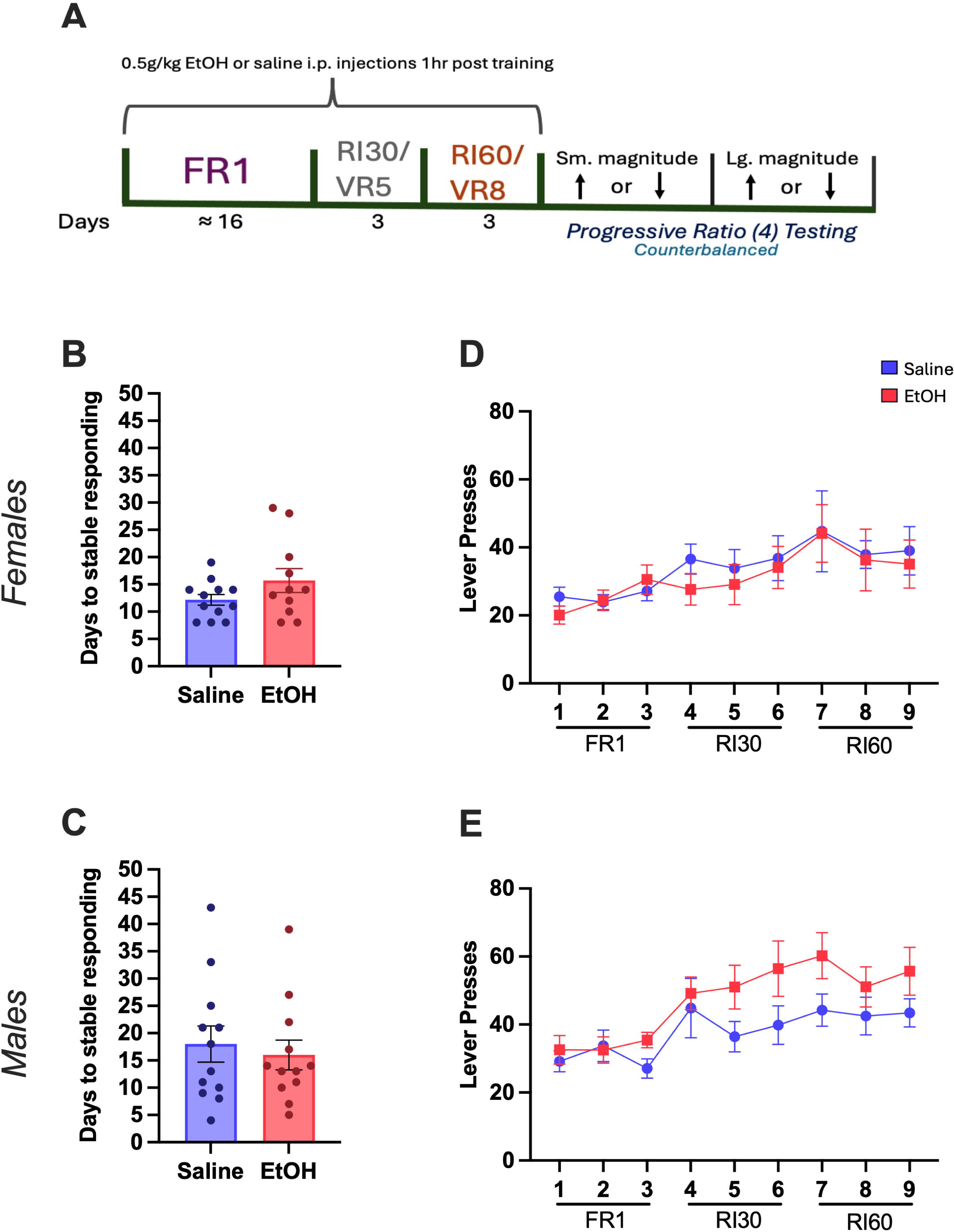
Low-dose ethanol did not impact acquisition. (**A**) Timeline of behavioral experiment. (**B,C**) Low-dose ethanol exposure did not alter the number of days required to achieve stable responding on an FR1 schedule in either female (**B**) or male (**C**) mice. (**D, E**) Low-dose ethanol exposure did not alter number of lever presses across training schedules in female (**D**) or male (**E**) mice. Females-saline: n = 12, ethanol: m = 11; males-saline: n = 12, ethanol: n = 12. Data represent mean +/− SEM.

### Chronic low-dose ethanol exposure alters sucrose reward motivation in female – but not male - mice when changes in reward magnitude occur during progressive ratio testing

To investigate the effects of chronic, low-dose ethanol on reward motivation in response to changes in reward magnitude, changes in breakpoints during PR testing were assessed. Because this task was self-paced, the number of training sessions and ethanol injections varied between animals (13-39 sessions). The number of sessions was not predictive of performance in the PR tests in either females [Pearson’s correlation: Days of EtOH exposure x Breakpoints: *lg. reduction*: R^2^ = 0.03839, p = 0.5637; *sm. reduction*: R^2^ = 0.002531, p = 0.8832; *sm. increase*: R^2^ = 0.01578, p = 0.7128; *lg. increase*: R^2^ = 0.1884, p = 0.1823] or males [Pearson’s correlation: Days of EtOH exposure x Breakpoints: *lg. reduction*: R^2^ = 0.03434, p = 0.5642; *sm. reduction*: R^2^ = 0.003731, p = 0.8504; *sm. increase*: R^2^ = 0.002578, p = 0.8755; *lg. increase*: R^2^ = 0.05750, p = 0.4529]. Control and low-dose ethanol-exposed mice exhibited similar overall mean breakpoints in both males and females following either a small reduction [*females:* t = 0.7273, df = 21, p = 0.4750; **Fig 2A**; *males:* t = 1.807, df = 22, p = 0.0845; **Fig 2B**] or a large reduction [*females:* t = 0.7309, df = 21, p = 0.4729; **Fig 2C**; *males*: t = 0.6052, df = 22, p = 0.5513; **Fig 2D**]. Investigation of individual differences showed that low-dose ethanol exposure shifted behavioral response patterns in female mice. Specifically, an observed shift in the distribution of female mice exhibiting changes in behavior was observed, where a greater proportion of ethanol-exposed female mice changed their response patterns following a small reduction in reward magnitude [X^2^(2, 23) = 6.378, p = 0.0412; **Fig 2A**]. This was not observed in males [X^2^ (2, 24) = 2.762, p = 0.2514; **Fig 2B**]. Following a large reduction in reward magnitude, no differences in breakpoints between control and low-dose ethanol-exposed females [X^2^ (2, 23) = 2.071, p = 0.3551, **Fig 2C**] or low-dose ethanol-exposed male mice were observed [X^2^ (2, 24) = 0.2338, p = 0.8897; **Fig 2D**].

**Figure 2:**
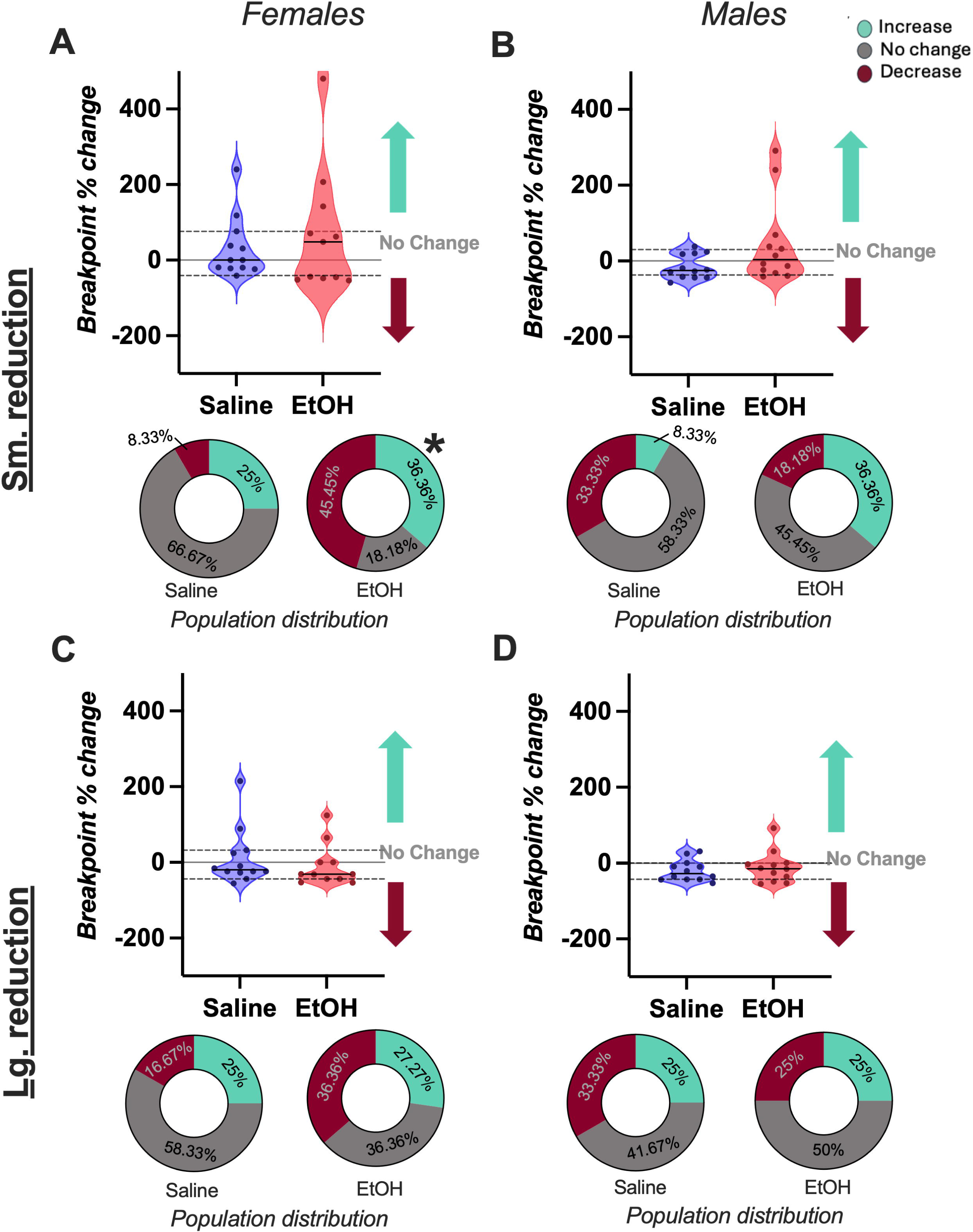
Low-dose ethanol altered sensitivity to small reductions in reward magnitude in female mice. (**A**) Female mice exposed to low-dose ethanol exhibited divergent responding when a small reduction in reward magnitude occurred. (**B**) Low-dose ethanol exposure had no effect on responding in male mice when there was a small reduction in reward magnitude. (**C**, **D**) When there was a large reduction in reward magnitude, there was no effect of ethanol in either female or male mice. X^2^, *p < 0.05. Females-saline: n = 12, ethanol: n = 11; males-saline n = 12, ethanol: n = 12, dashed line represent upper and lower quartiles with solid line representing mean.

When there was a small increase in reward magnitude, low-dose ethanol-exposed female mice exhibited significantly higher overall mean breakpoints compared to control female mice [unpaired t-test, t = 2.103, df = 21, p = 0.0477; **Fig 3A**] this effect was not observed in male mice [t = 1.060, df = 22, p = 0.3006; **Fig 3B**]. When there was a large increase in reward magnitude, both female and male mice responded similarly to controls [females: t = 0.4723, df = 21, p = 0.6416; **Fig 3C**; males; t = 0.2374, df = 22, p = 0.8145; **Fig 3D**]. When there was small increase in reward magnitude, both female [(X^2^ (2, 23) = 2.644, p = 0.2666, **Fig 3A**] and male mice [X^2^ (2, 24) = 1.333, p = 0.5134, **Fig 3B**] did not show significant differences in distribution in responding. Similarly, when there was a large increase in reward magnitude, both females [(X^2^ (2, 23) = 3.448, p = 0.1784, **Fig 3C**] and male [X^2^ (2, 24) = 1.333, p = 0.5134, **Fig 3D**] mice did not exhibit significant differences in population distribution versus controls. These findings indicted that the effects of low-dose ethanol on sensitivity to changes in reward were restricted to small changes (reduction or increase) as no effects were observed when there were large changes (reduction or increase).

**Figure 3:**
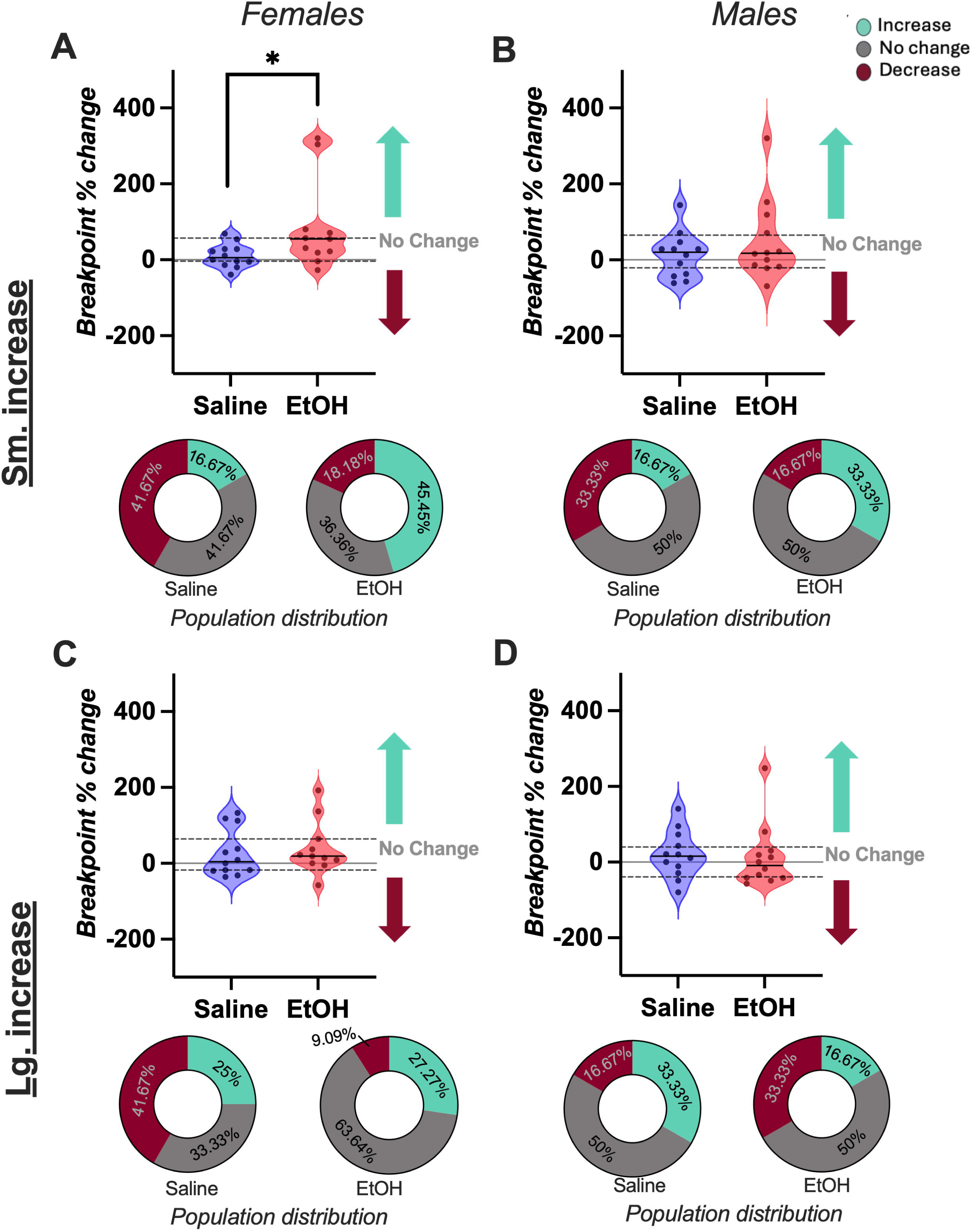
Low-dose ethanol altered responding in female mice when there was a small increase in reward magnitude. (**A**) Ethanol-exposed female mice increased responding when there was a small increase in reward magnitude in comparison to saline female mice. (**B**) Low-dose ethanol did not alter male mice responding when there was a small increase in reward magnitude. (**C**, **D**) Low-dose ethanol did not alter responding in female or male mice when there was a large increase in reward magnitude. *p < 0.05. Females-saline: n = 12, ethanol: n = 11; males-saline n = 12, ethanol: n = 12, dashed line represent upper and lower quartiles with solid line representing mean.

### Low-dose ethanol exposure alters behavioral strategy in a sex-specific manner

To probe the effects of chronic low-dose ethanol exposure on underlying behavioral strategies during PR testing, we quantified “checking” behavior as defined by percent of lever presses followed by a magazine entry for the entire PR session as well as within session as each PR block progressed.

When there was a small reduction in reward magnitude, we observed a significant decrease in overall checking behavior during the PR session [unpaired t-test: t = 2.168, df = 21, p = 0.0418; **Fig 4A (top)**] as well as within session in ethanol-exposed female mice in comparison to controls [two-way rmANOVA: *PR block*: F(2.383,26.21) = 1.323, p = 0.2863, Greenhouse-Geisser corrected; *EtOH*: F(1,21) = 5.438, p=0.0297; *PR block x EtOH*: F(10,110) = 0.5659, p = 0.8385; **Fig 4A (bottom)**]. This main effect of treatment was not observed in males when there was a small reduction in reward magnitude [t = 0.08621, df = 22, p = 0.9321, **Fig 4B (top)**; *PR block*: F(2.041,29.13) = 13.89, p<0.0001, Greenhouse-Geisser corrected; *EtOH*: F(1,22) = 0.2465, p = 0.6245; *PR block x EtOH*: F(11,157) = 0.3515, p = 0.9720; **Fig 4B (bottom)**]. There was an effect of PR block in checking behavior as the session progressed with a significant decrease in checking behavior during blocks with lever requirement 9 to 45 in comparison to the 1^st^ PR block [Dunnett’s multiple comparison: all p’s < 0.0259; **Fig 4B (bottom)**].

**Figure 4:**
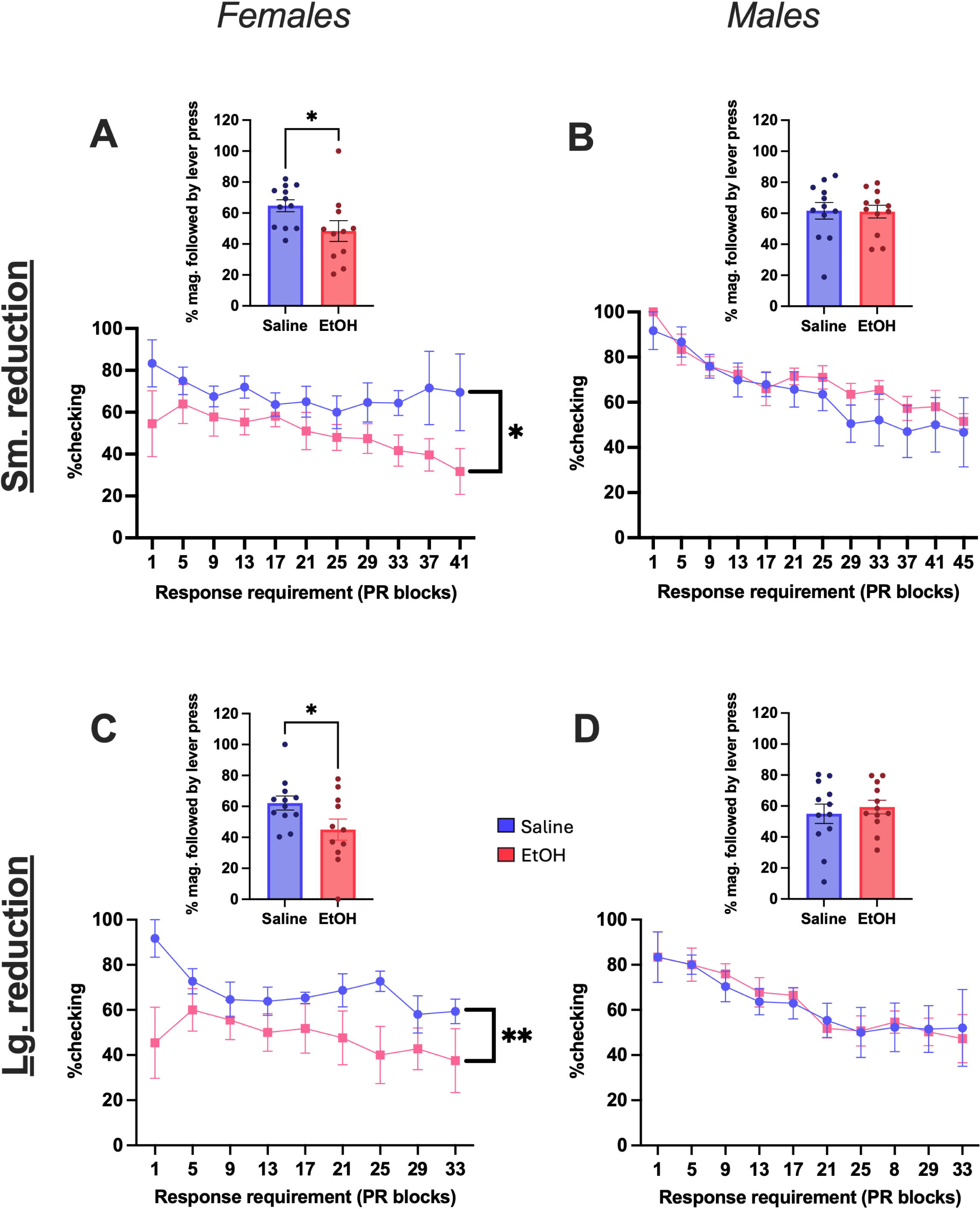
Analysis of ‘checking’ behavior during progressive ratio (PR) 4 testing when there was a reduction in reward magnitude. (**A**)Ethanol-exposed female mice reduced percent checking behavior during total (top) and within (bottom) PR testing session when there was a small reduction in reward magnitude. (**B**) Low-dose ethanol exposure did not affect percent checking behavior in male mice for total session or within session analysis when there was a small reduction in reward magnitude. (**C**) Ethanol-exposed female mice reduced percent checking behavior in total session (top) and within (bottom) PR testing session analysis when there was a large reduction in reward magnitude. (**D**) There was no effect of low-dose ethanol exposure on checking behavior in male mice, for the total or within PR4 testing session analysis, when there was a large reduction in reward magnitude. **p < 0.01, *p < 0.05. Females-saline: n = 12, ethanol: m = 11; males-saline: n = 12, ethanol: n = 12. Data represent mean +/− SEM.

Chronic low-dose female mice had lower checking behavior than controls when there was a large reduction in reward magnitude. This was observed both in overall [t = 2.121, df = 21, p = 0.0460; **Fig 4C (top)**] and within session analysis [*PR block*: F(2.900, 36.25) = 1.765, p = 0.1726, Greenhouse-Geisser corrected; *EtOH*: F(1, 21) = 8.279, p = 0.0090; *PR block x EtOH*: F(8, 100) = 1.162, p = 0.3299; **Fig 4C (bottom)**]. There was no effect of ethanol exposure on overall percent magazine checking [t = 0.5698, df = 22, p=0.5746; **Fig 4D (top)**] or within PR session analysis in mice male when there was a large reduction in reward magnitude [*PR block*: F(2.624, 38.19) = 4.873, p = 0.0077, Greenhouse-Geisser corrected; *EtOH*: F(1, 22) = 0.1189, p = 0.7335; *PR block x EtOH*: F(9, 131) = 0.3501, p = 0.9560; **Fig 4D (bottom)**]. There was a significant decrease in checking behavior, regardless of treatment, during the block in which the lever requirement was 25 in comparison to the 1^st^ PR block of the session [p = 0.0290; **Fig 4D (bottom)**].

When there was a small increase in reward magnitude, there was a significant reduction in checking behavior by ethanol-exposed female mice in overall PR session [t = 2.471, df = 21, p = 0.0221; **Fig 5A (top)**] but not when within session checking behavior was analyzed to that of control female mice checking behavior [*PR block*: F(2.109, 31.19) = 14.41, p<0.0001, Greenhouse-Geisser corrected; ETOH: F(1, 21) = 3.453, p = 0.0772; *PR block x EtOH*: F(14, 207) = 1.405, p = 0.1528; **Fig 5A (bottom)**]. We did observe an effect of PR block with a decrease in checking behavior in blocks with a lever requirement of 13 to 57 in comparison to the 1^st^ block regardless of treatment [all p < 0.0088; **Fig 5A (bottom)**]. There was no effect of low-dose ethanol on male mice overall checking behavior [t = 0.05214, df = 22, p = 0.9589; **Fig 5B (top)**] or within PR session analysis when there was a small increase in reward magnitude [*PR block*: F(5.024, 74.70) = 16.21, p<0.001, Greenhouse-Geisser corrected; *EtOH*: F(1, 22) = 1.540, p = 0.2277; *PR block x EtOH*: F(15, 223) = 0.5917, p = 0.8799; **Fig 5B (bottom)**]. We did observe an effect of PR block with a decrease in checking behavior during blocks with a lever requirement of 9 to 61 in comparison to the 1^st^ PR block [all p’s < 0.0029, **Fig 5B (bottom)**].

**Figure 5:**
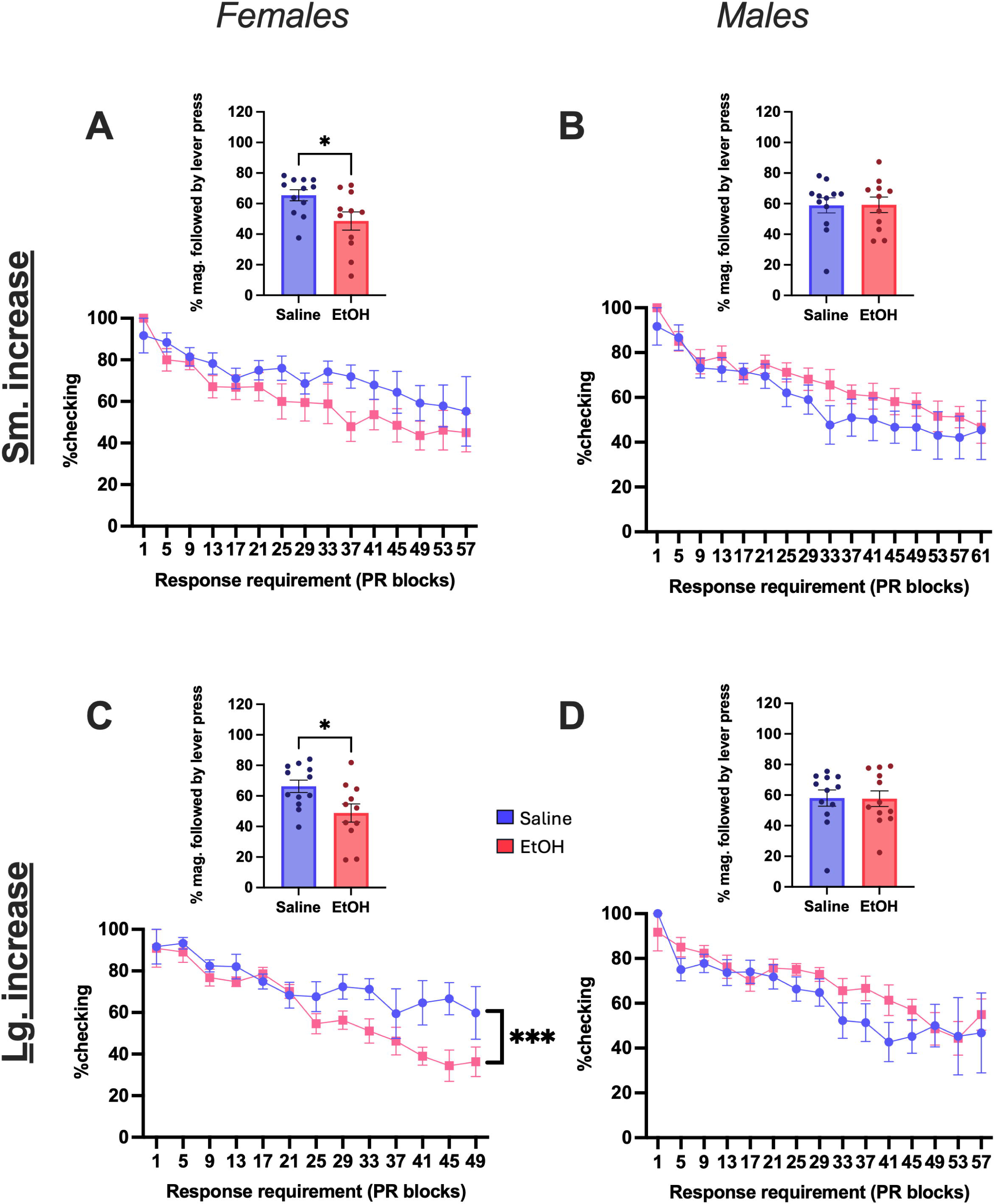
Analysis of ‘checking’ behavior during progressive ratio (PR) 4 testing when an increase in reward magnitude occurred. (**A**) Ethanol-exposed female mice reduced percent checking behavior in total (top) but not within PR testing session analysis when there was a small increase in reward magnitude. (**B**) Low-dose ethanol did not impact percent checking behavior in male mice in total session or within PR4 testing session analysis when there was a small increase in reward magnitude. (**C**) Ethanol-exposed female mice reduced percent checking behavior for total (top) and within (bottom) PR testing session analysis when there was a large increase in reward magnitude. (**D**) Low-dose ethanol exposure did not impact checking behavior in male mice, in total or within PR4 session analysis, when there was a large reduction in reward magnitude. ***p < 0.001, *p < 0.01. Females-saline: n = 12, ethanol: m = 11; males-saline: n = 12, ethanol: n = 12. Data represent mean +/− SEM.

When there was a large increase in reward magnitude, there was a significant decrease in checking behavior in both overall session analysis [t = 2.476, df = 21, p = 0.0219; **Fig 5C (top)**] as well as within PR session analysis in ethanol-exposed female mice [*PR block*: F(3.456, 52.99) = 13.61, p < 0.0001, Greenhouse-Geisser corrected; *EtOH*: F(1, 21) = 7.804, p = 0.0109; *PR block x EtOH*: 12, 184) = 1.681, p = 0.0738; **Fig 5C (bottom)**]. There was an effect of PR block with reduced checking behavior in blocks with a lever requirement of 25,33,37,41,45 and 49 vs the 1^st^ PR block of the session in female mice [all p’s < 0.047; **Fig 5C (bottom)**]. In response to a large increase in reward magnitude, there was no difference in percent ‘checking’ during overall session [t = 0.06169, df = 22, p = 0.9514, **Fig 5D (top)**] or within PR session analysis in ethanol male mice in comparison to controls [*PR block*: 4.003, 59.76) = 17.76, p < 0.0001, Greenhouse-Geisser corrected; *EtOH*: F(1, 22) = 0.05818, p = 0.8116; *PR block x EtOH*: F(14, 209) = 1.106, p = 0.3542; **Fig 5D (bottom)**]. There was a significant decrease in checking behavior in male mice during blocks with a lever requirement of 9 to 57 vs the 1^st^ PR block of the session [all p’s < 0.0380; **Fig 5D (bottom)**].

### Low-dose ethanol alters reward-circuit engagement following reward seeking

To investigate sex-specific effects of low-dose ethanol on neural substrates involved with reward motivation, we analyzed colocalization of a retrograde tracer injected within the NAc and cFos activity in NAc-projecting regions following sucrose reward seeking (15-minute RI60 session) in male and female mice who had a history of exposure to either saline or ethanol (**Fig 6A**). The percent of NAc-projecting neurons from the BLA, prelimbic cortex (PrL) or infralimbic cortex (IL) which expressed cFos (% cFos+ and GFP+/total GFP+), the percent of all cFos expressing neurons that projected to the NAc from the BLA, PrL or IL (% cFos+ and GFP+/total cFos+), and overall BLA, PrL and IL cFos+ cells were quantified.

**Figure 6:**
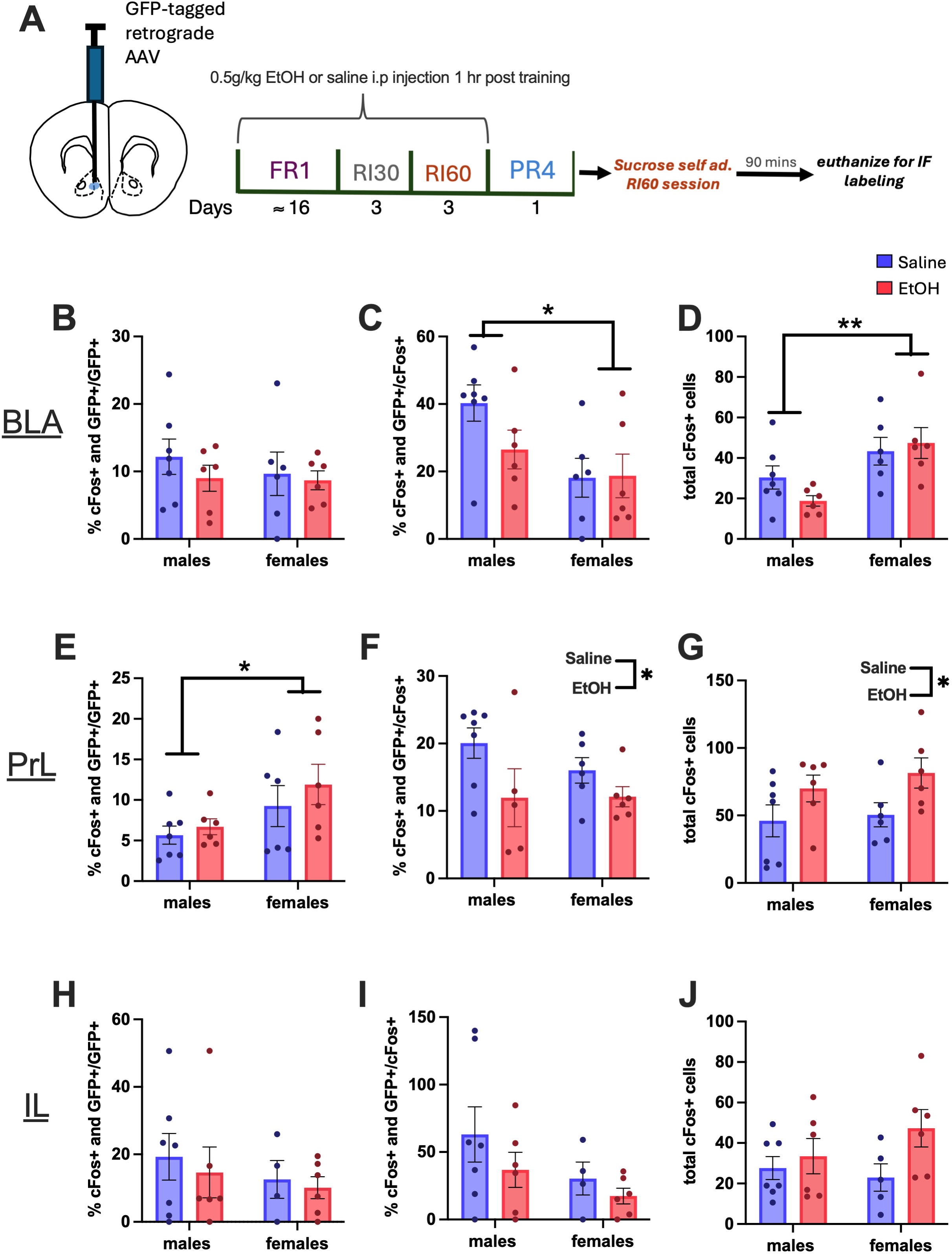
Low-dose ethanol had brain-region specific effects on neural activity. (**A**) Timeline of stereotaxic surgery, chronic low-dose ethanol exposure during behavioral training and sacrifice post reward seeking session for cFos immunofluorescence staining. (**B)** Neither a history of low-dose ethanol exposure nor sex altered percent of BLA projections to the NAC expressing cFos. (**C**) The proportion of cFos expressing cells projecting to the NAc within the BLA was greater in males than females, with no effect of ethanol exposure. (**D**) Female mice had a greater total of cells within the BLA expressing cFos than male mice, with no effect of ethanol exposure. (**E)** Female mice showed greater percent of PrL cells projecting to the NAc expressing cFos than males. (**F)** Ethanol-exposed mice showed reduction in PrL cells expressing cFos which projected to the NAc to that of controls. (**G**) Ethanol-exposed mice had a greater number of cells within the PrL expressing cFos than controls. Neither a history of low-dose ethanol exposure nor sex altered percent of IL projecting cells to the NAc expressing cFos (**H**) percent of cFos expressing cells which project to the NAc within the IL (**I**) or total cells within the IL expressing cFos (**J**). **p < 0.01, *p < 0.05. Females-saline: n = 6, ethanol: n = 6; males-saline: n = 6, ethanol: n = 6, Data represent +/− SEM.

Viral tracer surgery took place prior to ethanol exposure. We normalized co-expressing and cFos+ cells to each animal’s count of baseline GFP+ cells to account for any variation in AAV uptake. No significant difference in GFP+ cells between female and male mice were observed within the BLA [unpaired t-test: t = 0.05852, df = 21.28, p = 0.9539], PrL [t = 02.034, df = 15.54, p = 0.0594] or IL [t = 0.7773, df = 21.98, p = 0.4453]. Within the BLA, there was no effect of sex or treatment on percent of BLA projections to the NAc that expressed cFos [two-way ANOVA; *Sex:* F(1,21) = 0.3363, p = 0.5681; *EtOH*: F(1,21) = 0.7320, p = 0.4019; *Sex x EtOH*: F(1,21) = 0.2044, p = 0.6558; **Fig 6B**]. However, females exhibited a lower percentage of cFos expressing cells that projected from the BLA to NAc than males [two-way ANOVA; *Sex*: F(1,21) = 6.561, p = 0.0182]. There was no effect of ethanol treatment or interaction [*EtOH*: F(1,21) = 1.268, p = 0.2728; *Sex x EtOH*: F(1,21) = 1.500, p = 0.2342; **Fig 6C**]. This stands in contrast to greater total cFos expression in the BLA in females than males [two-way ANOVA: *Sex*: F(1,21) = 11.97, p=0.0023], potentially indicating activity in another BLA circuit in females [*EtOH*: F(1,21) = 0.3957, p=0.5361; *Sex x EtOH*: F(1,21) = 1.694, p = 0.2072; **Fig 6D**].

Within the PrL, there was a main effect of sex such that a greater percentage of PrL projections to NAc expressed cFos in female mice as compared to male mice [two-way ANOVA: *Sex*: F(1,21) = 5.521, p = 0.0286; **Fig 6E**]. There was no effect of ethanol treatment or interaction [*EtOH*: F(1,21)=0.9713, p=0.3356; *Sex x EtOH*: F(1,21) = 0.1868, p = 0.6700; **Fig 6E**]. In contrast, for both female and males there was a main effect of ethanol such that ethanol-exposed mice exhibited a lower percentage of cFos expressing cells that projected from the PrL to the NAc [two-way ANOVA: *EtOH*: F(1,21) = 5.636, p = 0.0277; *Sex x EtOH*: F(1,21) = 0.6829, p = 0.4184; **Fig 6F**] but a greater total of PrL cFos expressing cells when compared to saline controls [two-way ANOVA: *EtOH*: F(1,21) = 6.622, p = 0.0177; *Sex x EtOH*: F(1,21) = 0.7450, p = 0.7450; **Fig 2G**]. There was no effect of sex on cFos expressing cells projecting from the PrL to the NAc [two-way ANOVA: *Sex*: F(1,21) = 0.5879, p = 0.4522; **Fig 6F**] or total cFos expression within the PrL [two-way ANOVA: *Sex*: F(1,21) = 0.5569, p = 0.4638; **Fig 6G**].

Within the IL, there was no effect of sex or ethanol treatment on circuit-specific or total cFos expression [percent of IL projections to the NAc projections that expressed cFos: two-way ANOVA: *Sex*: F(1,21) = 0.7489, p = 0.3976; *EtOH*: F(1,21) = 0.2997, p = 0.5905; *Sex x EtOH*: F(1,21) = 0.02830, p = 0.8682; **Fig 6H**; percent total cFos that project to NAc: two-way ANOVA: *Sex*: F(1,21) = 2.794, p = 0.1110; *EtOH*: F(1,21) = 1.573, p = 0.2250; *Sex x EtOH*: F(1,21) = 0.1807, p = 0.6755; **Fig 6I**; total cFos: two-way ANOVA: *Sex*: F(1,21) = 0.3470, p = 0.5624; *EtOH*: F(1,21) = 3.785, p = 0.0659; *Sex x EtOH*: F(1,21) =1.412, p = 0.2487; **Fig 6J**]. Overall, these findings show that the effects of low-dose ethanol on NAc-projecting circuit engagement during reward seeking depend on sex and exposure and differ across cortical and subcortical regions.

## Discussion

There is increasing evidence of neurobiological and behavioral consequences of repeated or chronic exposure to lower doses of ethanol, though this remains understudied. Previous findings have identified alterations in motivated reward seeking and limbic circuitry (Bryant et al., 2022, 2023, 2024; Corbit et al., 2012; Cui and Koob, 2017; Fillmore and Rush, 2001). To further characterize low-dose ethanol effects on reward seeking and neural correlates, the current experiments investigated the consequences of chronic, low-dose ethanol exposure on motivated reward seeking following changes in reward magnitude. While no effects of low-dose ethanol exposure on acquisition of sucrose self-administration were observed, low-dose ethanol had sex-specific effects on behavioral response to change in reward magnitude, such that ethanol-exposed females – but not males – exhibited greater behavioral shifts following changes in reward magnitude compared to ethanol-naive mice. This was accompanied by a change in response strategy, with reduced reward magazine ‘checking behavior’ in ethanol-exposed females, but not males, during self-administration.

The absence of any effects of low-dose ethanol exposure on the acquisition of sucrose self-administration replicates our previous findings using a similar ethanol exposure procedure (Bryant et al., 2022, 2023, 2024) and further aligns with findings that higher ethanol exposure using vapor inhalation also does not impact overall sucrose self-administration (Barker et al., 2020; Renteria et al., 2018). Despite the similar acquisition curves and overall response rates between ethanol-exposed and control mice, the neural circuits engaged by reward seeking – as indicated by cFos expression – were impacted by sex and/or a history of ethanol exposure, with differences observed in the basolateral amygdala and prelimbic cortex. Consistent with our findings, cortico-limbic structures are particularly vulnerable to chronic ethanol exposure (Abernathy et al., 2010; Gass and Chandler, 2013; Klenowski et al., 2021; Koob and Volkow, 2016; Pleil et al., 2015; Shields et al., 2022; Stuber et al., 2011), with sex-specific effects of higher dose ethanol exposure on cFos expression within NAc circuitry (Chan et al., 2025).

The BLA, a key structure within the mammalian reward circuit, is thought to encode reward value and contribute to flexible actions (Ambroggi et al., 2008; Keefer et al., 2022; Mahler and Berridge, 2012; O’Doherty, 2003; Wassum, 2022). Activity within the BLA during reward seeking has been associated with ‘updating’ changes in reward value with disruptions in BLA activity corresponding to impaired ability to update behavior, potentially producing maladaptive reward seeking (Hinz et al., 2025; Jezzini and Padoa-Schioppa, 2020; Keefer et al., 2022; Wassum, 2022; Wassum et al., 2012). BLA’s function associated with the ability to update behavior consistent with reward value may be mediated by BLA projections to the NAc (Mahler and Berridge, 2012; Stuber et al., 2011; Wassum, 2022; Wassum and Izquierdo, 2015; Wellman et al., 2005). In both non-human primates and rodents, disconnection of the BLA and NAc impaired sensitivity to outcome devaluation, implicating this circuit in the ability to use reward value to guide behavior (Ambroggi et al., 2008; Wellman et al., 2005).

The current findings identified sex differences in cFos expression following self-administration in the BLA, independent of ethanol exposure. Despite overall higher cFos counts in the BLA of female mice compared to males, cFos expression was lower in BLA projections to the NAc in females. These findings not only identify greater activity of BLA neurons following self-administration in females, it also may suggest that BLA circuits other than the projection to the NAc are recruited during self-administration in female mice. This may be consistent with other findings of sex differences in BLA and NAc excitability and plasticity, with females exhibiting greater excitability in BLA (Price and McCool, 2022, 2021). This may impact circuit recruitment as well as behavioral strategies in response to changes in reward magnitude in a sex-specific manner (Copenhaver and LeGates, 2024; Price and McCool, 2022, 2021). While chronic low-dose ethanol effects on BLA cFos were not observed, sex differences in BLA cFos activity following self-administration may reflect differential sensitivity of female mice to changes in reward value. However, the current studies only assessed cFos expression following a standard self-administration session. It is possible that probing cFos expression following a session in which reward magnitude was altered would have revealed effects of chronic low-dose ethanol.

In contrast to findings in the BLA, where sex but not ethanol exposure history impacted cFos expression, cFos counts in the PrL were mediated by both ethanol exposure and sex. Following self-administration, male and female mice had similar total cFos expression within the PrL, however, female mice exhibited greater cFos expression than males within PrL to NAc projections, regardless of ethanol treatment. Total cFos expression was increased in the PrL of mice both male and female mice with a history of chronic, low-dose ethanol exposure. Despite this, the percentage of PrL to NAc projections expressing cFos was reduced in ethanol-exposed mice, suggesting that the increase overall PrL increase in ethanol-exposed mice was driven by PrL projections to other targets. One function of the PrL is evaluation of motivational conflicts or contexts that influence whether goal-directed actions should be suppressed or initiated (Green and Bouton, 2021; Shipman et al., 2018) thus, the PrL is essential for mediating flexible behavior and reward-driven decision making (Killcross and Coutureau, 2003; Wassum et al., 2012; Woon et al., 2020). While the IL is also implicated in the regulation of reward seeking, no changes in IL cFos were observed in this paradigm, suggesting that these findings do not generalize across the medial PFC. While the current study assesses the neural activation for nondrug self-administration, others have reported distinctions in the ensembles engaged by ethanol and sucrose reward (Ostlund et al., 2022) and drug/nondrug rewards (Pfarr et. al., 2018) within the medial PFC, which may have important implication for ethanol effect on ethanol or drug self-administration. Though not quantified in the current manuscript, the PrL also projects to the BLA. These projections are also involved in the regulation of reward seeking and may contribute to low-dose ethanol effects on behavior (Cardinal et al., 2002; Land et al., 2014; Song et al., 2020; Tavares et al., 2023). These changes in PrL activity following chronic low-dose ethanol exposure may independently impact the ability to flexibly regulate behavior or may drive deficits through interactions with downstream structures like the BLA. Mechanistic studies selectively modulating these structures and circuits will provide a more complete causal perspective on the contribution of these circuits to low-dose ethanol-induced behavioral alterations.

Reward motivation is determined by the value of the reward and the effort (cost) in seeking and obtaining the reward (Albert-Lyons et al., 2024; Bissonette and Roesch, 2016; Roesch and Olson, 2007). The current findings suggest that even at lower doses, chronic ethanol may impact the flexible relationship between effort and reward magnitude. Following either a large increase or decrease in reward magnitude, no effect of chronic, low-dose ethanol on reward motivation in either male or female mice in comparison to controls was observed. However, when changes in reward magnitude in either direction (decreases or increases) were smaller, ethanol-exposed female mice exhibited greater changes in behavior compared to controls. Following a small reduction in reward magnitude, low-dose ethanol female mice exhibited divergent behavioral responses. One population of female mice increased responding for the smaller reward, while another population reduced responding, suggesting that while ethanol exposure increased behavioral sensitivity to small reward reductions in female mice, this resulted in distinct behavioral phenotypes. In clinical studies, women have been reported to work harder for less reward, displaying higher motivation in the pursuit of goals or reward to that of men (Lewis et al., 2023). The current data suggest that at lower doses, ethanol may promote one similar phenotype in which a subset of female mice were willing to work harder for less reward. These divergent behavioral phenotypes observed in female mice may reflect vulnerabilities induced by chronic, low-dose ethanol exposure. For example, failure to persist in seeking goals when reward value is reduced may indicate risk for depression-related behaviors (Admon and Pizzagalli, 2015; Vrieze et al., 2013). Future work should consider whether these divergent populations reflect risk for neuropsychiatric illness, such as depression, or subsequent development of AUD.

In contrast to findings following reward reductions, low-dose ethanol-exposed female mice showed a greater increase in reward motivation following a small increase in reward magnitude in comparison to control female mice. This effect was not observed in chronic, low-dose ethanol-exposed male mice. Notably, previous work has found an overall increase in reward motivation as measured on the PR in low-dose ethanol-exposed male, but not female, mice (Bryant et al., 2023). Together, this suggests that chronic, low-dose ethanol exposure may alter sensitivity to changes in reward value in females, while having more general effects on reward motivation in males as observed in other higher dose ethanol exposure studies (Radke et al., 2017; Risher et al., 2013; Somkuwar et al., 2018). Though effects of ethanol on behavioral response were only observed following small changes in reward magnitude, ethanol-exposed female mice exhibited overall reduced ‘checking’ behavior – defined as percent of entries to the reward magazine that follow a lever press - across all PR test sessions. This decrease in ‘checking’ behavior appears to be a generalized behavioral outcome in chronic, low-dose ethanol exposure in female mice as we observed this altered behavioral microstructure across all testing conditions in ethanol-exposed female mice. This may be consistent with deficits in the use of action (lever press) outcome (reward delivery) contingencies to guide behavior in low-dose ethanol-exposed female mice. However, analysis of within-session checking behavior showed that chronic, low-dose ethanol female mice shifted checking behavior as the PR test sessions progressed, suggesting monitoring of reward contingencies. This would be consistent with previous work which found that female mice exposed to chronic, low-dose ethanol use checking behavior to guide their behavior in response to changes in contingency (Barker et al., 2024) and greater sensitivity to changes in outcome value in higher dose ethanol-exposed females (Barker et al., 2020; Pickens et al., 2020). Thus, the shift in behavioral strategy observed here during changes of reward magnitude may suggest increased reward sensitivity driven by increased goal-directed behavior in female mice. This is potentially consistent with findings that male, but not female, adult rats exhibit facilitated habit formation following higher doses of ethanol exposure (Barker et al., 2017), and when self-administering lower doses of ethanol (Barker et al., 2010). Together, this may suggest divergent effects of ethanol on value sensitivity and use of contingencies in males and females.

### Caveats and future considerations

The current study characterized the impacts of low-dose ethanol on non-drug reward seeking. It is possible that low-dose ethanol exposure would have differing effects on ethanol reward seeking as is observed following higher doses of ethanol. For example, previous work has shown that female mice subjected to chronic intermittent ethanol vapor inhalation increase reward sensitivity to ethanol (in the absence of conflict), promoting enhanced reward seeking in a conditioned place preference paradigm (Xie et al., 2019). It is also not clear how chronic, low-dose ethanol exposure would impact other effort or motivation assays other than the PR test used in this study. An important consideration to note in our current study is that ethanol exposure was administered by the experimenter in a bolus injection dose. While this procedure enabled controlled administration of this targeted dose of ethanol to determine effects on the brain and behavior in both males and females, and targeted timing of administration relative to behavior, it is likely that self-administered ethanol would engage different circuitry. This intraperitoneal injection method of delivery was selected to minimize stress of oral gavage, but this yields distinct metabolic impacts. In addition, while this dosing procedure yields similar BECs in male and female mice (Nothem et al., 2023), it is possible that the behavioral effects observed in females might emerge in males following an alternative dose of ethanol.

These studies assessed correlates of neural activation following reward seeking, cFos expression was used as a marker of putative cellular activity during reward seeking. This allowed investigation of NAc circuitry engagement following sucrose self-administration sessions, however, these experiments did not provide fine temporal resolution on changes in neural activity or characterize activity following changes in reward magnitude. It is important in future studies to probe changes in cellular activity across training to capture as these changes occur prior to testing. In this study sex and low-dose ethanol effects on PrL and BLA regional activity and their projections to NAc were identified. Future work should incorporate other circuitry such as PrL to BLA as well as BLA to PrL to help further identify the effects of low-dose ethanol on cortico-limbic structures involved with reward seeking and motivation. Incorporation of mechanistic focused studies would be beneficial to determine if the observed changes in substrate engagement and behavioral outputs were causally related.

### Conclusions

These findings indicate that repeated exposure to low-dose ethanol induced sex-specific effects, with ethanol-exposed female mice exhibiting alterations in motivated responding for sucrose and underlying behavioral strategies. This was accompanied by alterations in cortico-limbic circuit engagement following self-administration reward seeking, including low-dose ethanol effects on prelimbic activity in both male and female mice. This work highlights the importance of studying the effects of lower dose ethanol exposure on the brain and behavior to further understand how sex plays a role in the development of ethanol-induced aberrant reward seeking which may lead to increased risk of negative outcomes.

---

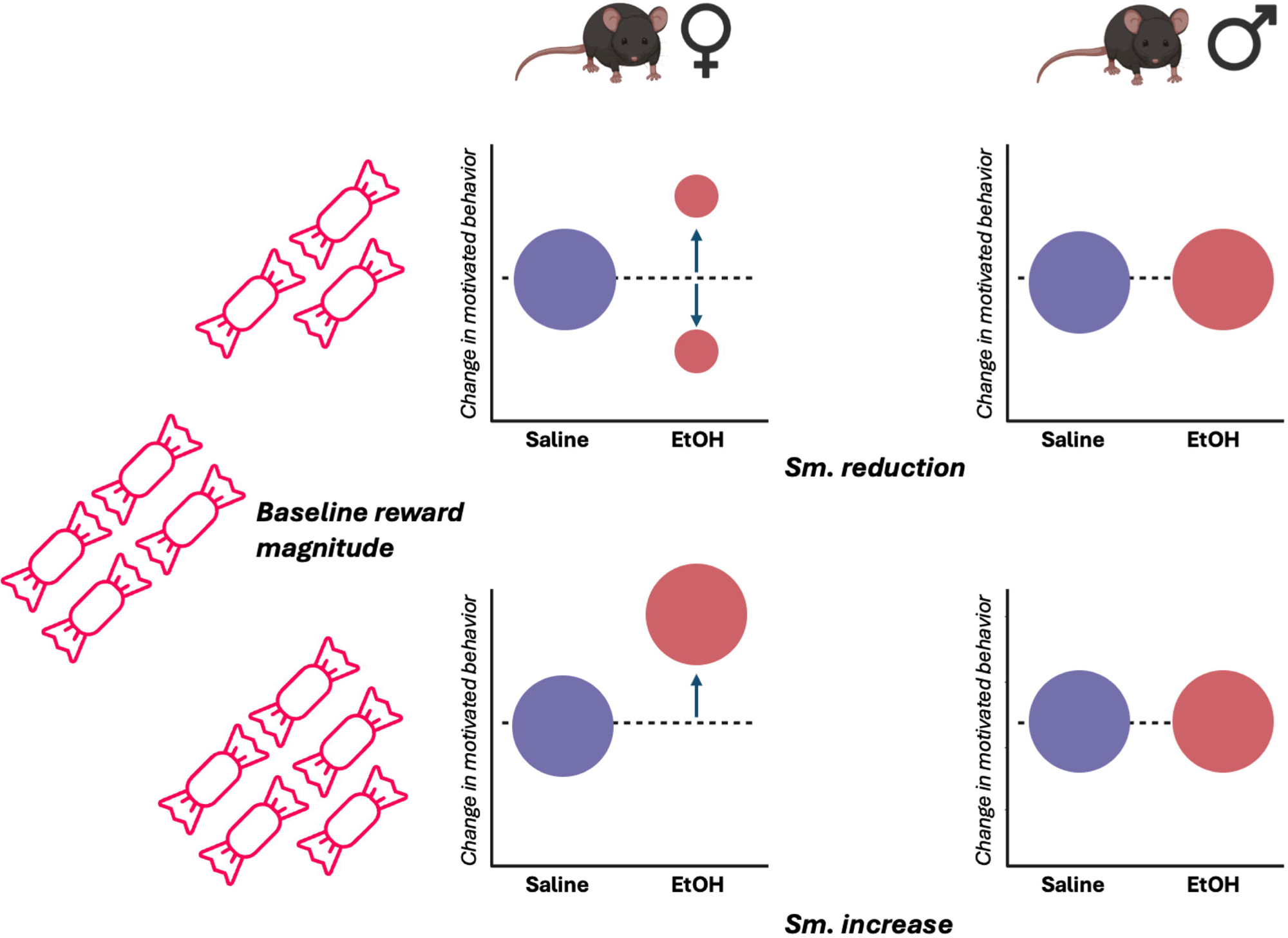

## Notes

### Competing Interest Statement

The authors have declared no competing interest.

### Summary of Updates

Figure 1 revised Figure order revised (Figure 2 now Figure 6 revised; Others moved 'up') Old Figure 3 revised (now Figure 2) Some text revisions for clarity of methods and rationale

